# A pathogenic CD4 T cell phenotype in experimental uveitis shares common features with other immune mediated inflammatory diseases

**DOI:** 10.1101/2025.06.14.659471

**Authors:** Amy Ward, Oliver H Bell, Luis Martinez-Robles, Phillipa J P Lait, Colin J Chu, David A Copland, Andrew D Dick, Lindsay B Nicholson

## Abstract

Murine ocular autoimmunity develops through 3 stages; prodrome, primary peak and secondary regulation. During the prodromal phase, leukocytes accumulate within the retina and vitreous. Using the adoptive transfer model of experimental autoimmune uveitis, we can analyse the disease course by tracking the transferred cells during disease initiation being recruited to the ocular environment throughout peak of disease to secondary regulation. During initiation (the prodrome) of disease ‘pathogenic’ transferred CD4^+^ T cells can be detected within the retina as well as an endogenous CD4^+^ infiltrate and as disease reaches peak, both transferred and endogenous CD4^+^ T cells can be found in large numbers in the retina. Active clinical disease resolves by day 21 but transferred CD4^+^ T cells persist within the retina when disease is in a clinically quiescent state. Concurrent transfer of RBP3 specific and OVA specific activated cells induces a similar clinical disease phenotype and time course. Both RBP3 and OVA specific cells are recruited during active clinical disease in equal measure showing that autoantigen specific CD4^+^ T cells induce susceptibility for recruitment of other activated CD4^+^ T cells. When analysing the endogenous and transferred CD4^+^ T cells by RNA sequencing, differences between the two sets of gene signatures highlight genes found in pathogenic T cells in other models, including upregulation of markers associated with cytokine interactions and NK cell mediated cytotoxicity. Due to the persistence of the original transferred population throughout clinical disease, in depth analysis of this population could suggest pathways contributing to ocular autoimmunity.

## 1. Introduction

Ocular autoimmune disease is one cause of non-infectious uveitis in humans, a group of conditions which as a whole lead to significant disability and burden on health-care [1]. In keeping with other immune mediated inflammatory diseases, it is associated with the development of autoantigen specific CD4 positive T cells [2, 3]. The ocular environment shares with several other tissues the property of immune privilege, characterised in health by tight endothelial barriers and limited immunosurveillance. In the healthy mouse eye this is reflected in the small number of lymphocytes that are recovered from the retina [4, 5]. In the animal model of ocular autoimmune disease, murine experimental autoimmune uveoretinitis (EAU), leukocyte accumulation in and around retinal vessels is a cardinal feature of disease, and work visualising cells using scanning laser ophthalmoscopy established that it is an early feature of EAU that can occur at a non-inflamed endothelial surfaces [6].

When ocular autoimmunity develops, there is a prodrome during which leukocytes accumulate in the retina. In the adoptive transfer model this occurs by day 2 after disease induction, and this is followed by an exponential increase in the number of cells recovered from the target organ, as the disease evolves from the prodromal stage to the initial peak [5]. Autoantigen specific CD4 positive T cells are critical drivers of this process and characterising the properties of these cells has been a focus of intense research over many years and in many different disease models [7–15]. Pro-inflammatory cytokines such as IFNγ and IL-17A/F are expressed by populations that are pathogenic, while cytokines such as IL-10 are crucial for the regulation and termination of immunity [16]. But other factors may contribute to pathogenicity, for example the expression of receptors associated with cytotoxicity [15], while other features may distinguish between subpopulations of CD4^+^ cells present in the infiltrate, for example changes in metabolic profile and the acquisition of stem like properties [13, 17].

To investigate the extended phenotype of pathogenic cells in EAU, we developed a model using a population of pathogenic CD4^+^ cells, transferred from donor mice with early EAU into naïve recipients. This allowed disease to develop in the absence of adjuvants. CD4^+^ cells were sufficient to initiate EAU and were characterised by an extended phenotype implicating functional properties beyond the production of pro-inflammatory cytokines. These cells set the scene for the accumulation of activated CD4^+^ cells within the immune privileged retinal environment.

## 2. Methods

### 2.1 Mice

OTII Ly5 mice CD45.1 (B6.CG-TG (TCRATCRB)425CBN/J) were obtained from JAX and bred at facilities at the University of Bristol. *CX3CR1^+/GFP^* mice on a C57BL/6 background were provided by Heping Xu (Queen’s University Belfast). Breeding colonies of homozygotes (confirmed by PCR) and offspring crossed with wildtype C57BL/6 mice were generated to give heterozygotes to run experiments assessing CX3CR1 expression.

OTII and CX3CR1^GFP/GFP^ mice were bred to create a OTII Ly5 CX3CR1^+/gfp^ mouse CD45.1^+^ CD45.2^+^ line. Blood was obtained from Ly5 OTII and Ly5 OTII CX3CR1^+/GFP^ for phenotyping using flow cytometry to check the transgenic TCR before experimentation. RAG2^−/−^ (B6.Cg-Thy1) mice were obtained from JAX and bred at facilities at the University of Bristol.

Mice were housed at the University of Bristol Animal Services Unit and housed under pathogen free conditions with water and food available *ad libitum*. All procedures were conducted in accordance with the United Kingdom Home Office and approved by the University of Bristol Ethical Review Group. Treatment of animals conformed to the Association for Research in Vision and Ophthalmology animal policy (ARVO statement for the use of animals in Ophthalmic and Vision Research).

### 2.2 Reagents

Murine RBP-3_629-643_ peptide (EAHYARPEIAQRARA) was synthesised by Severn Biotech (Worcestershire, UK) [18]. Peptide purity was determined by HPLC. Peptide was resuspended in water and stored as aliquots at −80°C.

### 2.3 Induction of EAU

#### Immunisation

Donor C57BL/6 CX3CR1^+/GFP^ mice were immunised subcutaneously in both flanks with a total of 40μg of RBP-3_629-643_ in phosphate buffered saline (PBS) emulsified (1:1 vol/vol) with Complete Freund’s Adjuvant (CFA) further supplemented with 1.5mg/ml *Mycobacterium tuberculosis* complete H37 Ra (BD Biosciences, Oxford UK). Donors also received 1μg of *Bordetella pertussis* toxin (Tocris, Bristol UK) given intraperitoneally (i.p).

#### RBP3 Cell Transfer

11 days after immunisation, spleen and lymph nodes were obtained from donors and seeded at 1-2×10^6^ cells per cm^3^ in 75cm^3^ flasks and cultured in complete medium (Dulbecco’s modified Eagle’s medium (DMEM) supplemented with 10% heat inactivated fetal calf serum (TCS Cellworks. UK), 100U/ml penicillin-streptomycin, and 2mmol/L L-glutamine (Invitrogen. Paisley, UK) supplemented with 10μg/ml RBP-3_629-643_ peptide and 10ng/ml IL-23 (R&D Abingdon, UK). 24 hours after cells are plated down the culture was further supplemented with 10ng/ml rIL-2 (Peprotech London, UK) in complete medium. At 72 hours, leukocytes were isolated by Ficoll density centrifugation and transferred by intraperitoneal injection, usually at 2×10^6^ total leukocytes per mouse in 100μl of PBS into naïve C57BL/6 recipients.

#### OVA Cell Transfer

Splenocytes from transgenic animals were stimulated in vitro for adoptive transfer for 72 hours. A naïve OTII spleen was prepared and seeded at 1-2×10^6^ cells per cm^3^ in 75cm^3^ flasks cultured in complete medium supplemented with 1mg/ml OVA peptide. After 72 hours, OVA reactive leukocytes were isolated using Ficoll density centrifugation and transferred by intraperitoneal injection into C57BL/6 recipients allelically marked by CD45.2. In some experiments, on the day of transfer recipients were anaesthetised using intraperitoneal injection of 90 μL/10 g body weight of a solution containing 6 mg/mL ketamine (Ketavet; Zoetis Ireland Ltd., Dublin, Ireland) and 2 mg/mL Xylazine (Rompun; Bayer plc, Newbury, UK) mixed with sterile water before intravitreally injecting 2µl of OVA peptide at 1µg/µl.

### 2.4 Clinical Assessment of EAU

At pre-defined experimental time points after disease induction mice were assessed clinically by using optical coherence tomography (OCT) (Days 2, 7, 14,21). Mice are firstly anaesthetised by intraperitoneal anaesthetic and pupils dilated by topical tropicamide 1% and phenylephrine 2.5%. Viscotears were used throughout imaging to lubricate the eye.

### 2.5 Isolation and flow cytometric analysis of retinal and vitreous infiltrate

Eyes were dissected by insertion into the limbal area and removal of retina and vitreous. Samples were dissociated by mechanical disruption before filtering through a 70 micron filter to acquire a single-cell suspension for analysis by flow cytometry.

Samples were incubated with purified rat anti-mouse CD16/32 Fc Block (Clone 2.4G2; BD Biosciences, Oxford UK) for 10 minutes at room temperature. Cells were then stained with fluorochrome-conjugated monoclonal antibodies against cell surface markers CD45.1, CD45.2, CD3e, CD4, CD8, CD11b, LY6G, LY6C (BioLegend, San Diego CA) at 4°C for 20 minutes.

Data was acquired using a 4-laser Fortessa X-20 flow cytometer (BD Cytometry Systems, Oxford UK). Data analysis was carried out using FlowJo software (Treestar, San Carlos California).

Absolute cell counts were calculated by known standard curve counts previously reported [4]. Briefly, a single cell suspension of splenocytes was generated and diluted to give a range of known concentrations of cells, all samples and standards were acquired in a 200µl total volume of FACS buffer at a fixed flow rate for 90 seconds. Based on data acquired from known standards a curve was generated to interpolate cell counts of ocular infiltrating cells from samples.

### 2.6 mRNA-Sequencing and analysis

Samples were prepared and sequenced using the SMART-Seq v4 Ultra Low Input RNA Kit for Sequencing as previously described [19]. Using the Galaxy platform, sequencing data was quality checked following adapter removal using FastQC. RNA STAR (2.7.8a, using mm10) was used to align reads and gene expression was quantified by featureCounts [20]. Differential gene expression analysis was carried out in R (4.3.1) using voom normalisation with sample specific weights in limma (3.56.2) [21, 22]; gene set enrichment was performed using fgsea (1.26.0).

### 2.7 Statistics

Data was analysed using GraphPad Prism 7 software (GraphPad Software Inc., San Diego CA). A non-parametric Kruskal -Wallis was used with a Kruskal-Wallis multiple comparisons test or a Man-Whitney test to generate statistical analysis of the data.

## 3. Results

### 3.1 RBP3 629-643 specific CD4^+^ T cells transfer uveitis

Pathogenic cells were harvested from donors 11 days after induction of the EAU model by immunisation with mouse RBP3 629-643 [18] and conditioned for adoptive transfer by culture in the presence of cognate antigen, IL-2 and IL-23 (Fig 1A). Prior to transfer, the CD45^+^ fraction included CD4^+^ cells (20%) CD8+ cells (10%) and CD11b+ cells (9%). The CD4^+^ cells had a mixed Th1 and Th17 profile, and following PMA/ionomycin stimulation, produced the signature cytokines IFNg and IL-17A at the same frequency (15% of the transferred CD4^+^ cells). By comparison, Foxp3 positive cells were only present in small numbers (1%) (Fig 1B). Leukocytes were transferred into wild type recipients across a range of concentrations (5×10^4^-5×10^6^) and disease was monitored using Topical Endoscopic Fundal Imaging (TEFI) and Optical Coherence Topography (OCT) to document incidence and progression [23, 24] (Fig 1C and Fig 1D). The adoptive transfer of 2×10^6^ leukocytes per recipient induced moderate disease in >80% of recipients that reached a peak by day 12. To investigate the pathogenic potential of the different lymphocyte cell types, 5×10^5^ CD4^+^ and CD8^+^ cells were FACS-sorted after conditioning (to a 99% pure population) and transferred into C57BL/6 RAG2^−/−^ mice. Severe clinical ocular disease was observed in recipients that received CD4^+^ cells but not CD8^+^ cells. This developed between day 14 and day 28, confirming that the CD4^+^ cells were necessary and sufficient to induce uveitis in the context of an intact innate immune cell population, whereas the CD8^+^ cells do not have this capacity (Fig 1E).

**Figure 1:**
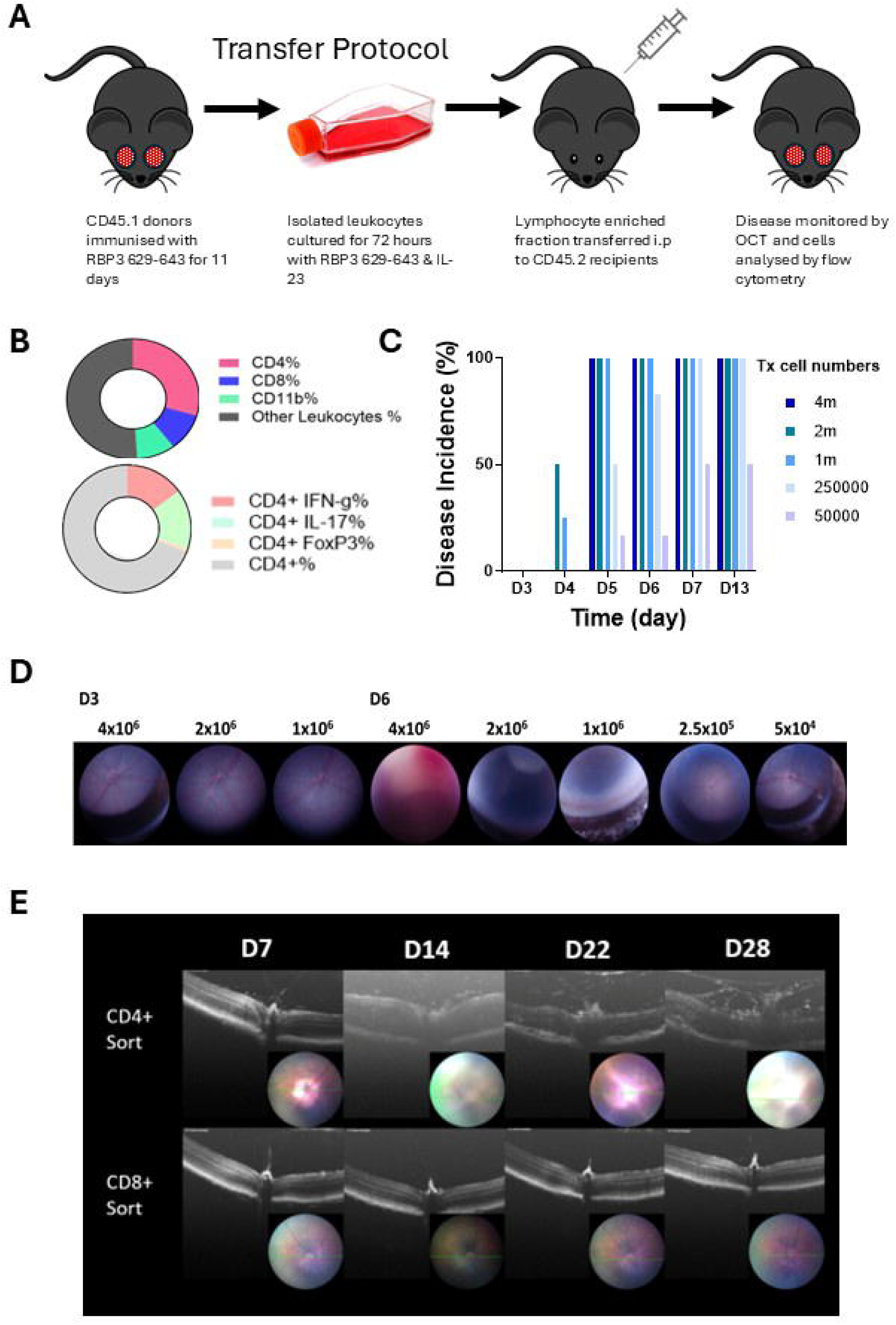
Adoptive transfer of uveitogenic leukocytes to induce EAU. (A) Experimental protocol overview for induction of EAU by adoptive transfer of uveitogenic leukocytes from CD45.2/CX3CR1^+/GFP^ into CD45.1 recipients. (B) Analysis of transferred CD45 positive leukocyte showing CD4 (20%), CD8 (10%) and CD11b (9%) and intracellular IFN-γ (8%), IL-17 (6%) and FoxP3 (1%). (C) To optimise leukocyte transfer to induce consistent disease phenotype, cell numbers were titrated and transferred into naïve recipients. Average disease scores were calculated using the method described in the methods section. (D) Topical Endoscopic Fundal Imaging (TEFI) and Optical Coherence Tomography (OCT) images were taken during the initiation of clinical disease and during peak clinical disease. (E) To test whether the CD4 transferred population is sufficient to initiate disease, sorted CD4+ and a CD8+ cells were transferred into RAG2^−/−^ mice and disease was monitored by OCT.

### 3.2 Early accumulation of ocular antigen specific cells following transfer

Leukocyte accumulation in the eye is a cardinal feature of uveitis and its animal model EAU. Previous work visualising cells in vessels by SLO [6] demonstrated that this can occur across a non-inflamed endothelium and studies in our laboratory has shown that the adjuvant components used in active immunisation lead early to a detectable increase in ocular CD4^+^ cell number, whether or not the stimuli are uveitogenic (Fig S1). The transfer of cells into naïve recipients, without the presence of adjuvant, provides a more sensitive approach than active immunisation to monitoring the progression of disease in vivo.

### 3.3 Transfer of antigen specific cells initiates an endogenous cell recruitment to the ocular environment

To investigate and compare pathogenic (ocular antigen specific and pro-inflammatory) versus non-specifically recruited CD4^+^ cells, we extended the transfer model (Fig. 1) by using allelically marked cells from donor mice, harvested 11 days after active immunisation and transferred in aliquots of 2×10^6^ total cells by intraperitoneal injection. Cells from the host (‘endogenous cells’) can then be distinguished from transferred cells (‘pathogenic cells’)using allelic markers.

Demonstrating the sensitivity of this approach to systemic perturbation, i.p. injection of tissue culture grade PBS alone led to a small but significant increase in ocular endogenous CD4^+^ cell recruitment (Fig 2A) which resolved to the level seen in unmanipulated mice over 7-14 days. The transfer of naïve TCR transgenic CD4^+^OT-II cells (OT-II-) or antigen activated and conditioned OT-II cells (OT-II+) did not alter this background effect on endogenous CD4^+^ cell recruitment, but when pathogenic autoantigen (RBP-3 629-643) reactive cells were used, endogenous CD4^+^ cell recruitment increased above the level seen following non-specific stimuli (Fig 2A). Naive OT-II-cells transferred into naïve mice were barely detectable in the eye 2 days after cell transfer of 2×10^6^ (Fig. 2B). Consistent with published work [6] when activated OT-II cells, conditioned using the same regimen as pathogenic uveitogenic cells were transferred, we can recover OT-II cells from the eye. Transferred pathogenic cells were retained at much higher levels at day 2, increasing further by day 7 as uveitis developed (Fig 2C and Table 1).

**Figure 2:**
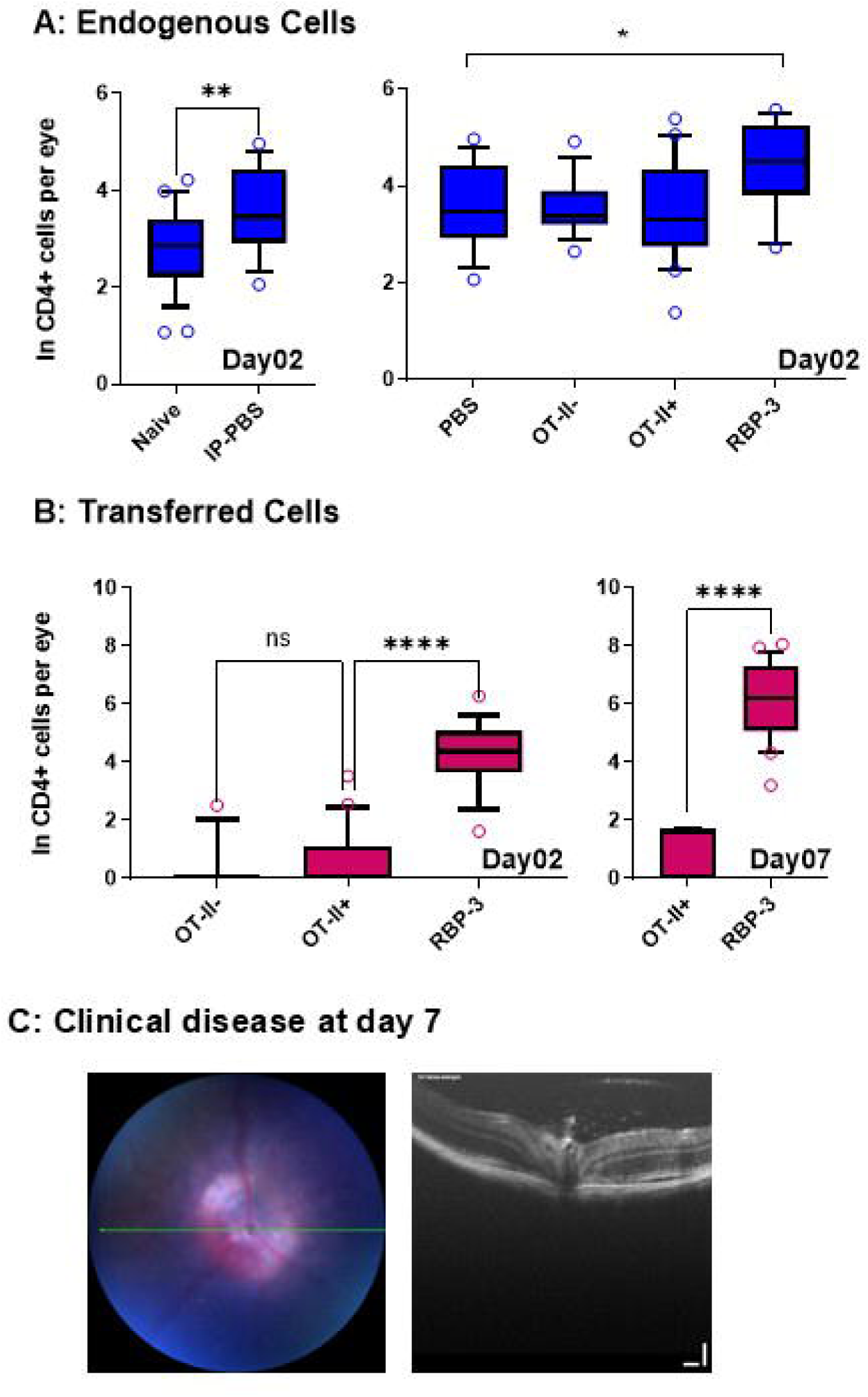
Quantification of endogenous and pathogenic ocular cellular infiltrate. (A) Endogenous cells were recovered at day 2 from the enucleated eyes of naïve animals or receiving aliquots of 2m transferred leukocytes i.p. and quantified by flow cytometry. (B) Transferred cells were recovered at day 2 or day 7 from the enucleated eyes of animals receiving aliquots of 2m cells i.p. Cell numbers are log transformed. Average absolute cell numbers: Panel A Naïve 16, IP-PBS 37, PBS 37, OT-II-36, OT-II+ 32, RBP-3 81. Panel B day02 OT-II-1, OT-II+ 2, RBP-3 74 day07 OT-II+ 2, RBP-3 431. (C) Representative clinical disease at day 7. Data was log transformed to normalise data. Comparisons of two-sample data by Mann-Whitney test, multiple comparisons using Kruskal-Wallis test carried out with GraphPad Prism software. * <0.05, ** <0.01 **** <0.0001

**Table 1.** Endogenous and transferred CD4 cell numbers recovered from the eyes of animals receiving PBS i.p., non-pathogenic cells (OT-II activated or unactivated transgenic T cells) or pathogenic (RBP3 629-643 specific cell line) cells. *p<0.05 compared with PBS or non-pathogenic endogenous cells at day 2; ****p<0.0001 compared with non-pathogenic transferred cells at day 2; ***p<0.005 compared with non-pathogenic transferred cells at day 2. **†**The transfer of pathogenic cells leads to a significant increase in all cell types on days 7 and 14 compared with day 2. p-values Kruskal-Wallis test with Dunn correction for multiple testing; p-adjust method Benjamini-Hochberg).

| CELL TYPE | TRANSFER | DAY |  |  |  |  |  |  |  |
| --- | --- | --- | --- | --- | --- | --- | --- | --- | --- |
|  |  | 2 |  | 7† |  | 14† |  | 21†† |  |
|  |  | Mean | Range (n) | Mean | Range (n) | Mean | Range (n) | Mean | Range (n) |
| ENDOGENOUS CD4 | PBS | 50 | 7-143 (16) |  |  |  |  |  |  |
|  | Non-pathogenic | 53 | 3-215 (50) |  |  |  |  |  |  |
|  | Pathogenic | 109* | 15-263 (18) | 3125 | 19-13320 (28) | 1128 | 18-4034 (27) | 128 | 8-891 (16) |
| TRANSFERRED CD4 | Non-pathogenic | 3 | 0-66 (50) |  |  |  |  |  |  |
|  | Pathogenic | 120**** | 5-534 (18) | 648 | 0-3165 (28) | 666 | 0-6727 (27) | 115 | 6-287 (16) |
| ENDOGENOUS CD8 | PBS | 39 | 10-154 (16) |  |  |  |  |  |  |
|  | Non-pathogenic | 29 | 0-127 (50) |  |  |  |  |  |  |
|  | Pathogenic | 62*** | 11-176 (18) | 1223 | 6-4034 (28) | 781 | 12-6129 (27) | 158<sup>s</sup> | 3-488 (16) |
| ENDOGENOUS CD11B | PBS | 1688 | 385-4866 (16) |  |  |  |  |  |  |
|  | Non-pathogenic | 2223 | 409-6009 (50) |  |  |  |  |  |  |
|  | Pathogenic | 1236* | 163-2750 (18) | 21743 | 658-73685 (28) | 1086 | 799-24042 (27) | 2083 | 340-8552 (18) |

### 3.4 Recruitment of non-RBP3 specific cells during active clinical EAU

By day 7 after cell transfer, clinical uveitis, detected by imaging of the retina and optical coherence tomography (OCT) is apparent in most animals and flow cytometry, to characterise the cellular infiltrate, reveals a mixture of endogenous and pathogenic leukocytes. We reasoned that the seven-day time span is sufficient for the development of an endogenous ocular antigen specific CD4^+^ T cell response, as well as for the induction of changes in the tissue to make it receptive to the retention of non-antigen specific cells. To test whether active uveitis produces an environment that is more retentive of non-antigen-specific CD4^+^ T cells, we analysed the recruitment of co-transferred activated TCR transgenic OT-II cells in the presence of EAU, seven days after cell transfer, compared with the transfer of OT-II cells alone. Using three different mouse strains, homozygous for CD45.1, homozygous for CD45.2 and double positive (CD45.1^+^CD45.2^+^), we could quantify three independent populations. Co-transfer of equal aliquots of pathogenic and activated non-ocular antigen specific OT-II T cells led to a large increase in the number of OT-II cells recovered from the eye at day 7 compared with experiments when only activated OT-II cells were transferred (mean 18 cells versus mean 2560; compare Fig. 2B and Fig. 3). The average CD4^+^ cell numbers of the different groups of cells at day 7 and day 14 were of similar magnitude (Day 7: endogenous 2625; OT-II 2560; pathogenic 1242. Day 14: endogenous 841; OT-II 404; pathogenic 305) (Table 2).

**Figure 3:**
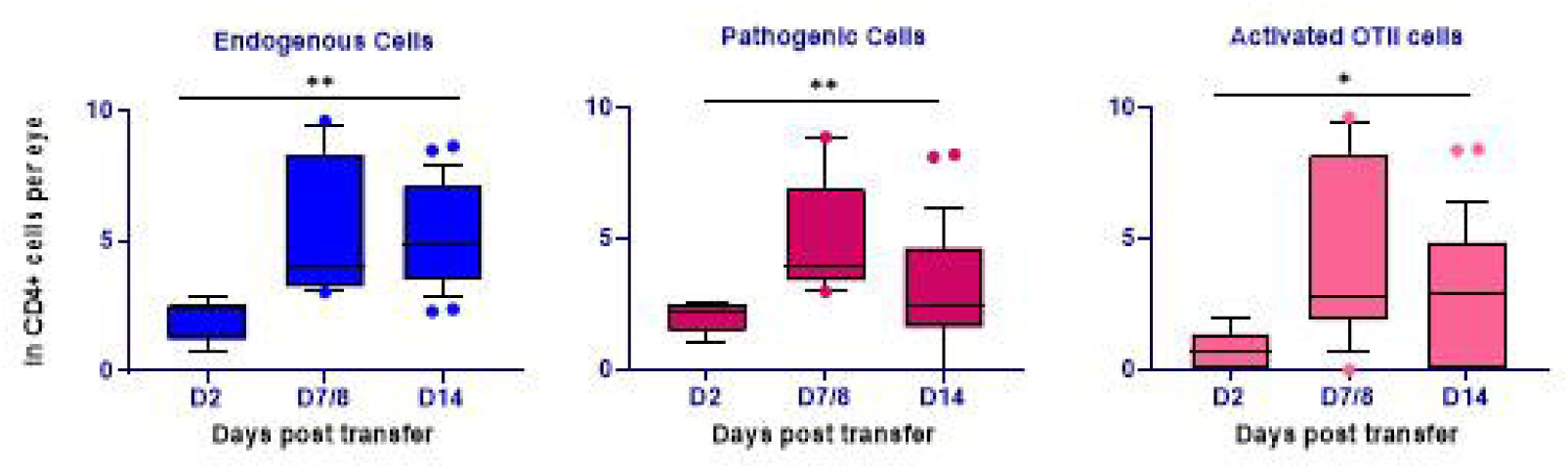
Quantification of endogenous, pathogenic and antigen non-specific cells recovered from eyes 2-14 days following co-transfer. Multiple comparisons assessed by Kruskal-Wallis test using GraphPad Prism software. * <0.05, ** <0.01.

**Table 2.** Pathogenic (RBP3 629-643) and non-pathogenic (OT-II TCR transgenic) were co-transferred at a ratio of 1:1. Cell infiltrate was quantified by flow cytometry.

CD4+ CELLS (± SE)
| DAY (N) | Pathogenic | Non-pathogenic | Endogenous |
| --- | --- | --- | --- |
| 7/8 (8) | 2144 ± 1096 | 4474 ± 2079 | 4573 ± 1999 |
| 14 (18) | 447 ± 402 | 598 ± 500 | 1222 ± 585 |

### 3.5 Local activation of antigen-specific CD4^+^ T cells leads to retention of activated CD4^+^ T cells within the ocular tissue

The co-transfer of pathogenic cells leads to a change in the capacity of ocular tissue to retain activated CD4^+^ T cells. These changes are tissue wide but one likely source of the signal that drives them is the local antigen specific activation of CD4^+^ T cells triggering the release of cytokines. To test this, we injected one eye (LE) with the cognate ligand for the CD4^+^OT-II T cells and the other eye (RE) with an irrelevant control peptide at the same time as transferring activated CD4^+^OT-II cells. This provides an activating antigen specific signal in only one eye and, as assessed by imaging, OCT and cell quantification, we found that clinical disease developed in this eye (LE) but not in the eye that received injection with a control irrelevant peptide (Fig 4).

**Figure 4:**
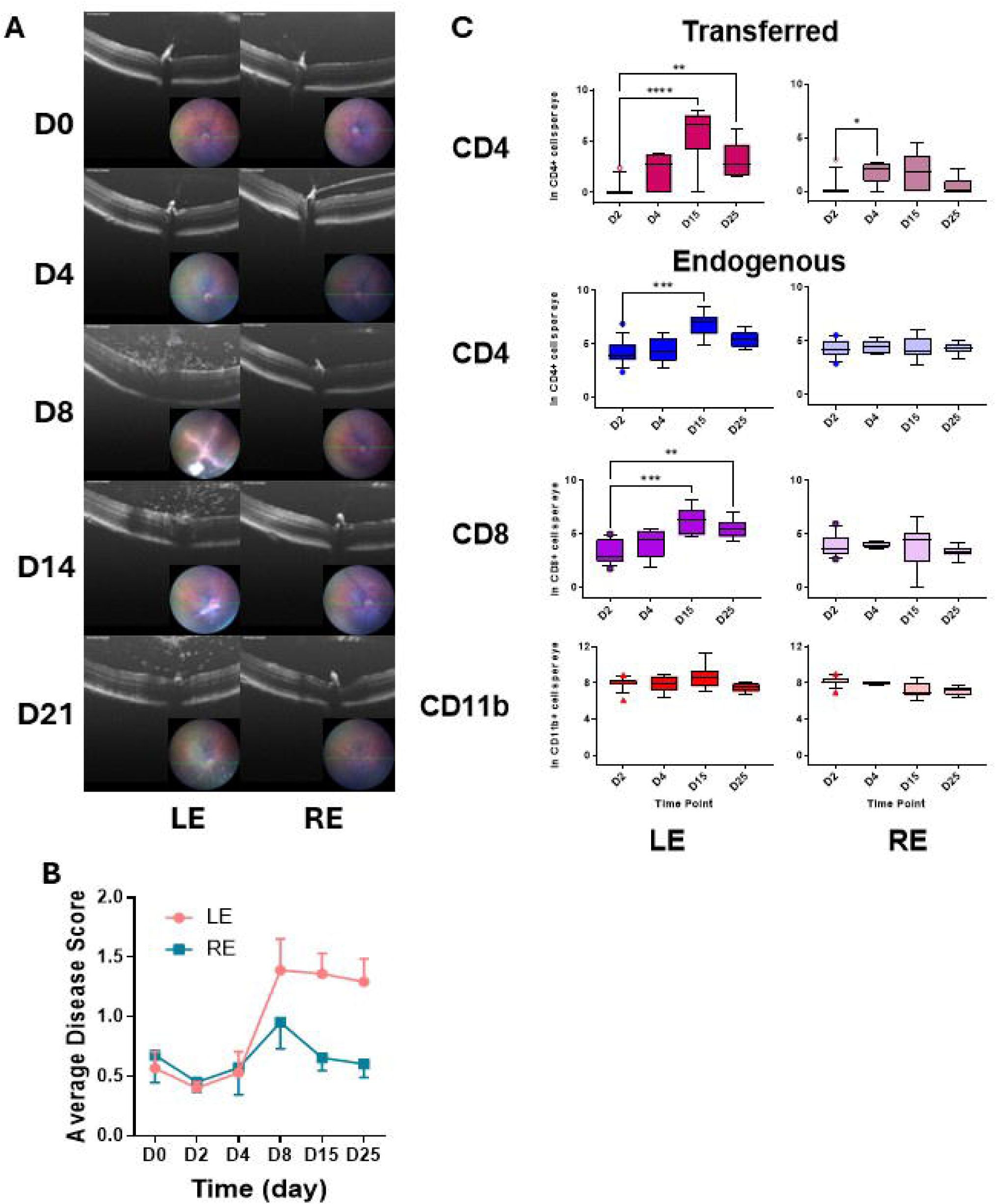
Local antigen presentation drives cell accumulation. (A) Clinical disease following cell transfer and intravitreal injection of OVA peptide (LE) of control peptide (RE). There is accumulation of cells reaching a peak around D14. (B) Blinded clinical scoring of disease in left and right eyes. (C) Transferred and endogenous cells accumulate in greater numbers in the eyes that received intravitreal injection of the cognate OVA peptide.

These experiments show that when activated in the eye, transferred pathogenic CD4^+^ cells are sufficient to establish an ocular microenvironment that is permissive for the retention of activated CD4^+^ T cells. When that antigen signal is localised to one eye, changes in the ocular environment are unilateral. We were then interested in tracking the long-term fate of these pathogenic pioneer cells by monitoring their presence as EAU evolved. We wished to understand whether over time the pioneering population was replaced by endogenous cells. We recorded clinical disease over 67 days, quantifying endogenous and pathogenic cells in the ocular infiltrate at different time points. We detected transferred CD4^+^ in the eye at all time points studied. These cells could also be detected at low levels at all time points in the spleen and lymph nodes (Mean 0.02% of total CD4^+^ cells in the spleen) (Fig 5).

**Figure 5:**
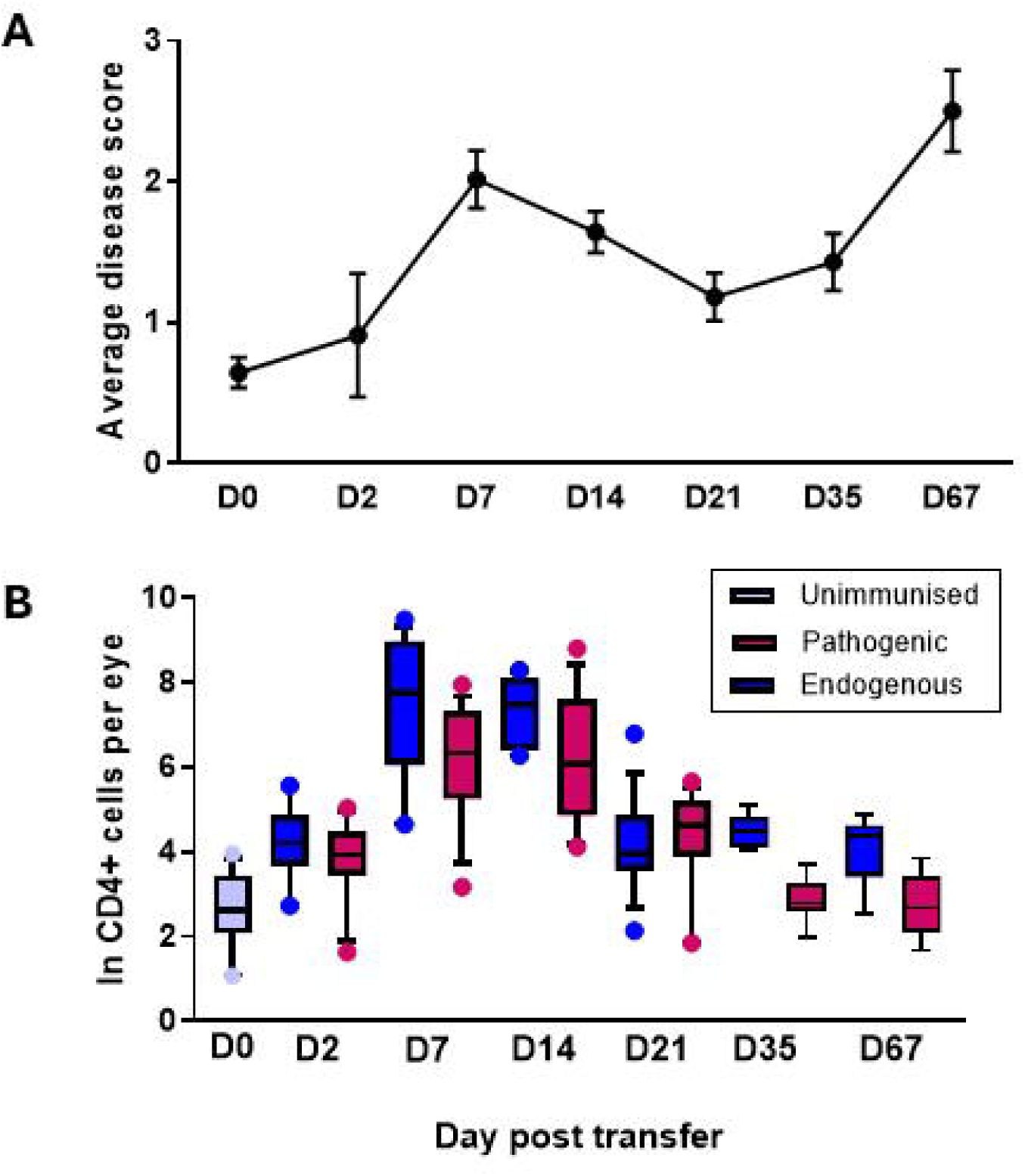
Clinical disease score varies of extended clinical follow-up (A). Pathogenic and endogenous cells can be recovered from retinas at all time points examined (B).

### 3.6 Pathogenic and endogenous CD4^+^ T cells have different gene signatures

The question then arose as to whether the local tissue microenvironment imprinted pathogenicity on lymphocytes or whether the differentiation of the pioneer CD4^+^ T cells played a more significant role in conferring pathogenic potential on different populations. Pathogenic and endogenous cells from the eye and spleen were analysed on D13 after transfer. We sorted 18-100 pathogenic or endogenous cells from individual eyes and 100 endogenous cells per sample from spleens of the same animals. T cells from the spleen had an activated (CD44^hi^, CD62L^lo^) phenotype. Using a validated protocol for low cell input RNA-Seq [19] and data processing in the open source Bioconductor framework, results were analysed first by Multidimensional scaling (MDS) plots of individual samples. These showed poor discrimination based on the tissue of origin (Fig 6A) but classifying cells as pathogenic or endogenous gave a good separation of samples and removing outliers yielded a set of data for further analysis which convincingly discriminated between transferred (13 samples) and pathogenic (11 samples) cells (Fig. 6B).

**Figure 6:**
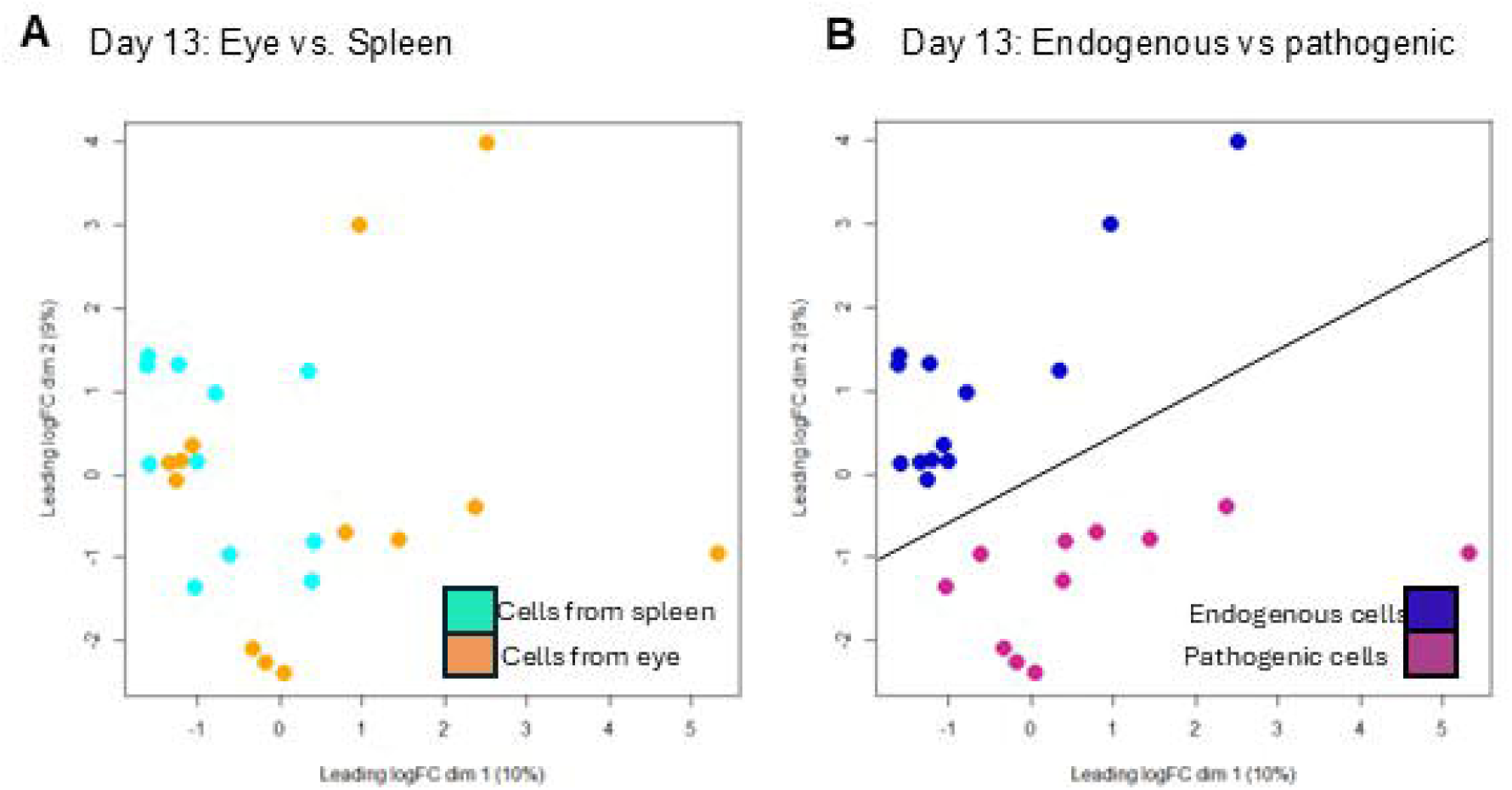
Following normalisation multidimensional scaling plots of CD4+ cells recovered from the eye or spleen show a mixture of phenotype (A). Classifying cells into pathogenic (transferred cells) or endogenous produces two distinct populations for differential gene expression analysis (B).

Following other analyses of lymphocyte signatures [13], genes that were differentially expressed with a fold change greater than 1.5 were identified. This revealed a total of 213 genes, 113 downregulated and 100 upregulated (Fig 7A). Genes of interest can be seen in a volcano plot (Fig 7B) and a heatmap (Fig. 7C). Clustering analysis of these genes revealed samples in pathogenic or endogenous groups and genes in 9 clusters, 4 with upregulated genes associated with pathogenic cells (Fig. S2) and 5 with genes that were downregulated in pathogenic cells (Fig. S3).

**Figure 7:**
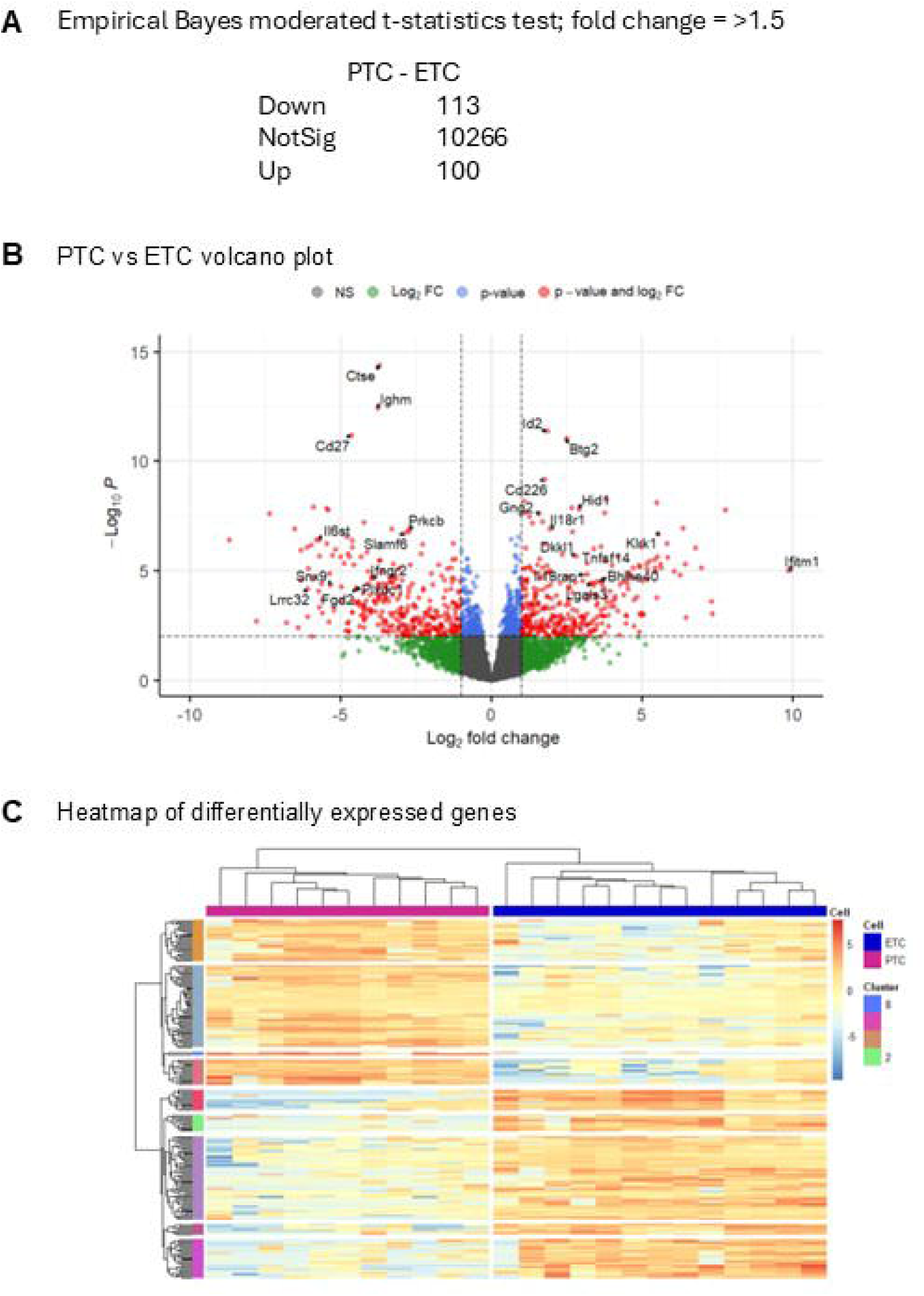
Differential gene expression (DGE) analysis. (A) With a threshold of fold change > 1.5, FDR 0.05 213 differential expressed genes were identified. (B) Volcano plot of DGE highlights immune relevant genes in up-regulated and down-regulated population. (C) Heat map of DGE with k-means clustering identifies 9 clusters.

To test the broader relevance of the pathogenic phenotype we carried out a gene-enrichment analysis using data from CD4^+^ cells harvested from the eyes of C57BL/6 animals with actively induced EAU. These cells were generated and analysed an independent experiment carried out in our laboratory. Cells were sampled close to the peak of disease, which in the actively induced model is 3-4 weeks post immunisation, here taken on day 26. Within a set genes from day 26, ranked by log fold change (logFC), the day 13 signature was significantly enriched (normalised enrichment score (NES) 1.69; adjP 8.18e-05) (Fig 8A).

**Figure 8:**
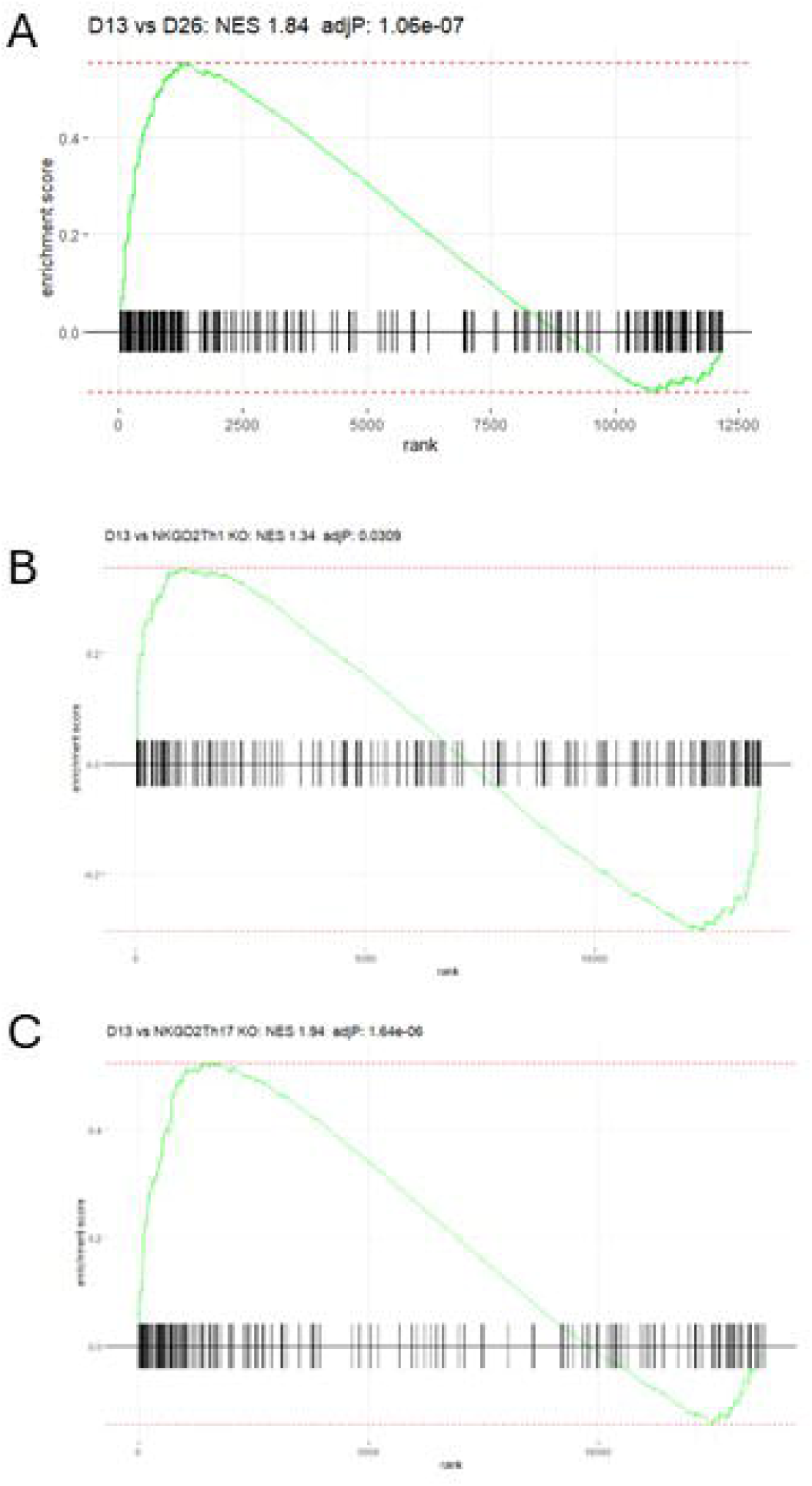
D13 signature found in active EAU and NKG2D knockouts. (A) The pathogenic gene expression signature (213 genes identified by DGE analysis is significantly enriched in CD4+ cells recovered from the retinas of animals with active EAU at D26. (B&C) The pathogenic gene expression signature is significantly enriched in NKG2DTh1 and Th17 cells compared with NKG2D KO cells with a higher enrichment score (1.94) in the Th17 analysis.

### 3.7 Pathogenic cells within the eye upregulate markers associated with cytokine interactions and NK cell mediated cytotoxicity

To better understand the pathological significance of the changes in gene expression seen in the pathogenic cell population we then carried out an enrichment analysis in the KEGG database which found that the 99 upregulated genes had a significant association with cytokine interactions and with NK cell mediated cytotoxicity. One significantly upregulated gene, *Klrk1* (coding for NKG2D) was of interest because its reported role in enforcing proinflammatory features on CD4^+^ T cells. We reanalysed data from this paper which compared the profile of wildtype or NKGD2D knockout cells differentiated to a Th1 or Th17 phenotype (GEO accession number GSE143522). We created gene lists ranked by logFC for differentially expressed genes in in vitro differentiated Th1 or Th17 cells from WT or NKG2D knockout donors. The NES was 1.4 and 1.85 for Th1 and Th17 cells respectively with a more significant association with Th17 cells (adjP 5.95e-05 vs 0.029). An examination of leading edge genes also implicated cytotoxicity as an important component of all these phenotypes (Supp Fig. 4). Finally, we considered relevance to human disease. We have previously characterised an immune signature associate with human inflammatory ocular disease [25]. Using the set of mouse orthologs of these human immune related genes that we have previously described (Supp T1) we found that the genes from this list were significantly enriched in the ranked day 13 gene set (NES: 1.73; AdjP 2.60e-05) supporting the relevance of the extended pathogenic phenotype in human disease.

## 4. Discussion

We developed the EAU transfer model, adapted from other models that have been successful applied to study EAE in C57BL/6 mice [26], using an ex-vivo expansion step that included recombinant IL-23 to produce a mixed population of Th1 and Th17 CD4^+^ cells. These cells produced uveitis by day 5 when 400,000 or more CD4^+^ cells, in an aliquot of 2×10^6^ cells were transferred (Fig. 1A-C). The pathogenicity of this cell population was likely enhanced by the presence of IL-23, which has recently been described as conferring a colitogenic potential on Th1 cells independent of its role in Th17 cell differentiation [27]. The pathogenic potential of these uveitogenic cells was localised within the CD4^+^ cell population, as shown by transfer into RAG knockout recipients (Fig.1 D,E), even though immunization with RBP3 629-643 does elicit autoantigen recognising CD8+ cells [18]. Using allelically marked donor cells allowed a very sensitive assay of changes in ocular immunosurveillance, assessed by FACS quantification of CD4^+^ cells recovered from retinas. Previous experiments have demonstrated that these numbers are unchanged following whole animal perfusion prior to FACS and represent lymphocytes that are either tissue resident or tightly associated with the vascular endothelium. In the transfer model we found that non-specific perturbation (injection of tissue culture grade PBS i.p.) had a small measurable effect on the pool of ocular resident CD4^+^ T cells (Fig. 2A). We reasoned that at least part of the explanation for this change likely arises from endothelial activation that increases the average dwell time of lymphocytes within the ocular microvasculature and demonstrates a very sensitive recalibration of immunosurveillance following non-specific stimuli. This interpretation is further supported by the demonstration that the introduction of ocular autoantigen specific cells leads to their accumulation within the retina and at the same time an increase in accumulation of endogenous cells above that induced by non-inflammatory signals (Fig. 2).

This autoantigen reactive CD4^+^ cell driven activation process effectively opens the tissue to the accumulation of activated CD4^+^ cells in much greater numbers than are seen in the eyes of unmanipulated animals. Because the antigen specificity of the recruited endogenous cells could not be determined, to test whether recruitment could be non-antigen specific, we transferred activated OT-II cells that have no known endogenous ligands in normal C57BL/6 mice and found that the evolving autoimmune inflammation efficiently recruits these cells to the retina as uveitis reaches its peak (Fig. 3). Carrying out this analysis at single timepoints shows comparable levels of transferred pathogenic cells and OT II TCR transgenic cells at the time of sampling. It remains unknown what the half-life of these different populations is within the eye. Prior work has shown that in EAU the ocular infiltrate is rapidly diminished by the administration of S1PR inhibitors [28] but also that certain subsets of lymphocytes are retained following this treatment [29] and there may be different classes of cells, with different dwell times within the tissue, that contribute to local immune memory [30].

To test if local antigen presentation is sufficient to support the retention of autopathogenic cells within the retina, we tested this by introducing the cognate peptide for the OT-II TCR transgenic T cells locally by intravitreal injection. Transferred OT-II cells were detected in both eyes following intravitreal injection but recruited CD4^+^ and CD8^+^ cells were found in significantly increased numbers only in the eye treated with the cognate peptide and not in the eye that received control peptide. This strongly supports the model in which local antigen presentation within the immune privileged tissue is critical in the disease process and is consistent with data in EAE pointing to a crucial local antigen presenting population [31] and also with recent data demonstrating the widely different potential of anatomically closely associated vascular beds to engage with the inflammatory process [32].

Following mice with EAU induced by cell transfer over an extended period of time provides data consistent with prior observations in actively induced EAU [18] that the immunosurveillance of the immune privileged retina is significantly reorganised over the long-term. The experiments reported here also allow us to conclude that the original T cells that instigated disease, or their clonal progeny, are retained. It is possible that these pioneer cells provide the source for a stem-like autoreactive pool, in a fashion that has been carefully established for CD8 cells in a model of type 1 diabetes [33] and identified at a transcriptomic level by extensive single-cell lymphocyte profiling under different inflammatory conditions [11]. Where in the EAU model these cells will be found is unknown, although the gut [34, 35] and the spleen [13] as well as draining lymph nodes and bone-marrow remain important candidate sites.

Following confirmation of the persistence of pioneer cells, one crucial question was the nature of their pathogenicity. We addressed this by using bulk RNA-seq of small (18-100) aliquots of cells from spleen and eye, defining the cells as pathogenic if derived from the transferred aliquot or endogenous if otherwise. Our initial analysis revealed that classifying cells by organ of origin produced fairly poorly discriminated populations compared with classification into pathogenic versus endogenous. This was a little surprising, in the context of literature that infers a strong effect of tissue origin on the phenotype of the recovered lymphocytes [13, 36, 37], but could reflect our sampling of an underlying dynamic process at this relatively early stage in disease, where cells are recirculating between the spleen and the ocular environment, especially as the cells sorted from the spleen were enriched for an activated/memory phenotype (CD44^hi^, CD62L^lo^).

We interrogated the extended phenotype of the pathogenic versus endogenous CD4^+^ T cells and identified 213 genes with significantly different levels of gene expression and a fold change of >= 1.5. Using clustering analysis (k-means) we divided the genes into 9 clusters. (Fig. 7, SF 2&3). We validated the relevance of this data set to uveitis by testing how the 213 significantly different genes related to a second independently derived RNA-Seq analysis of CD4^+^ cells carried out at day 26 on cells obtained from the eyes of mice with EAU induced by active immunisation. The significant enrichment (adjP 1.06e-07) of the pathogenic associated genes in the day 26 samples argues that this ‘signature’ is a common feature of multiple EAU models (Fig 8A). Extending this approach further, the study of NKG2D knockout T cells provided further evidence of the generality of the pathogenic signature in immune mediated inflammatory disease. When the NKG2D knockout cells were studied in experimental autoimmune encephalomyelitis, they were less pathogenic than their wild-type counterparts, supporting a role for this signature in influencing the level of tissue damage [15].

Many immune pathways are functionally conserved between animal models of autoimmunity and human disease. Within our ranked day 13 gene set genes that we have previously described as relevant in persistent ocular inflammation were significantly enriched (NES: 1.73; AdjP 2.60e-05). Validation in these different models raises the possibility that these pathways, that are driving ocular pathology, may also be important therapeutic targets in other immune mediated inflammatory diseases.

## 9. Acknowledgements

Acknowledgements- We would like to acknowledge the University of Bristol Flow cytometry facility: Dr Andrew Herman and Helen Rice and the University of Bristol Animal Services Unit.

