## Supplementary data for "A pathogenic CD4 T cell phenotype in experimental uveitis shares common features with other immune mediated inflammatory diseases"

### Slide 1
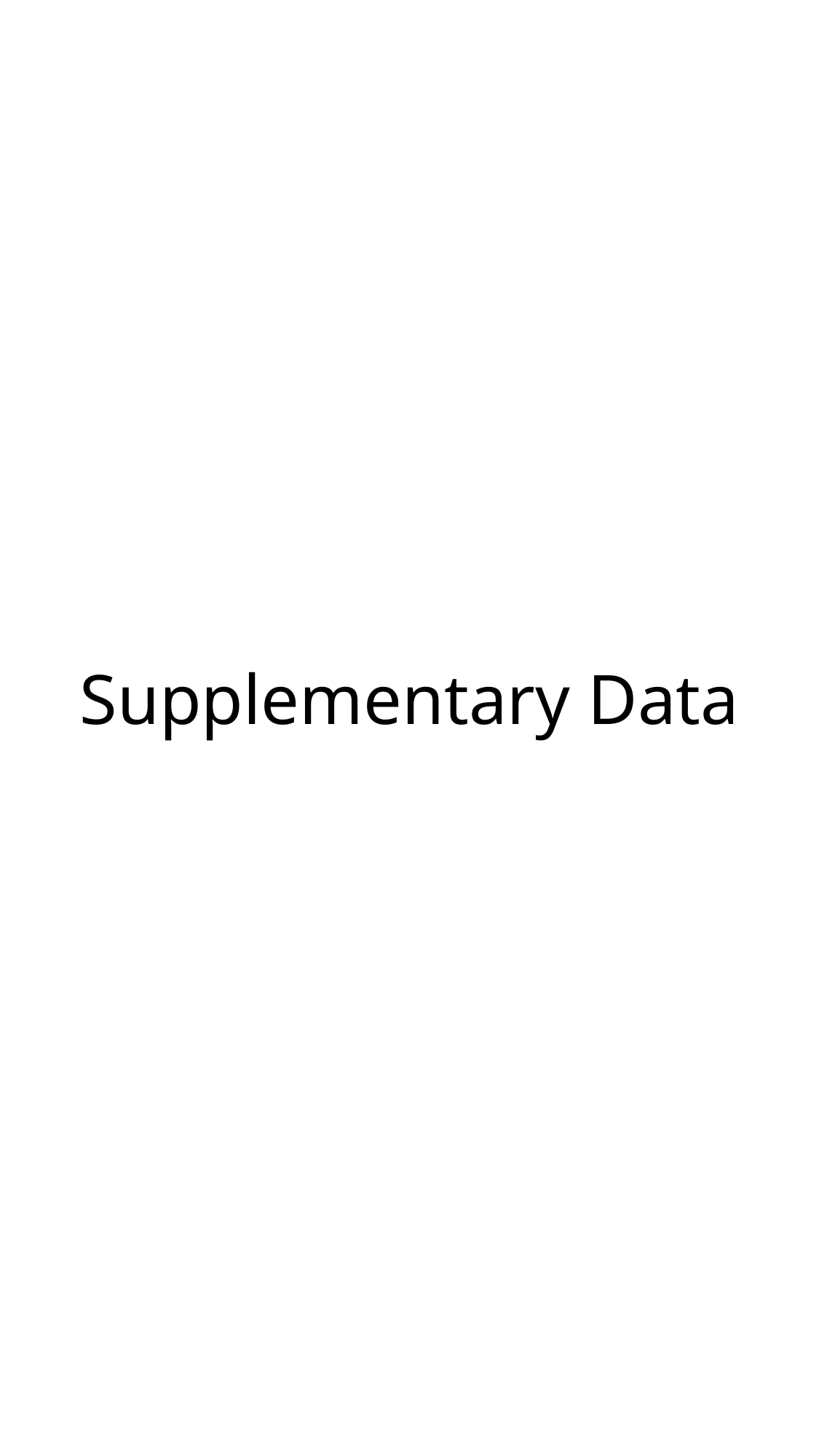

#### Supplementary Data

### Slide 2
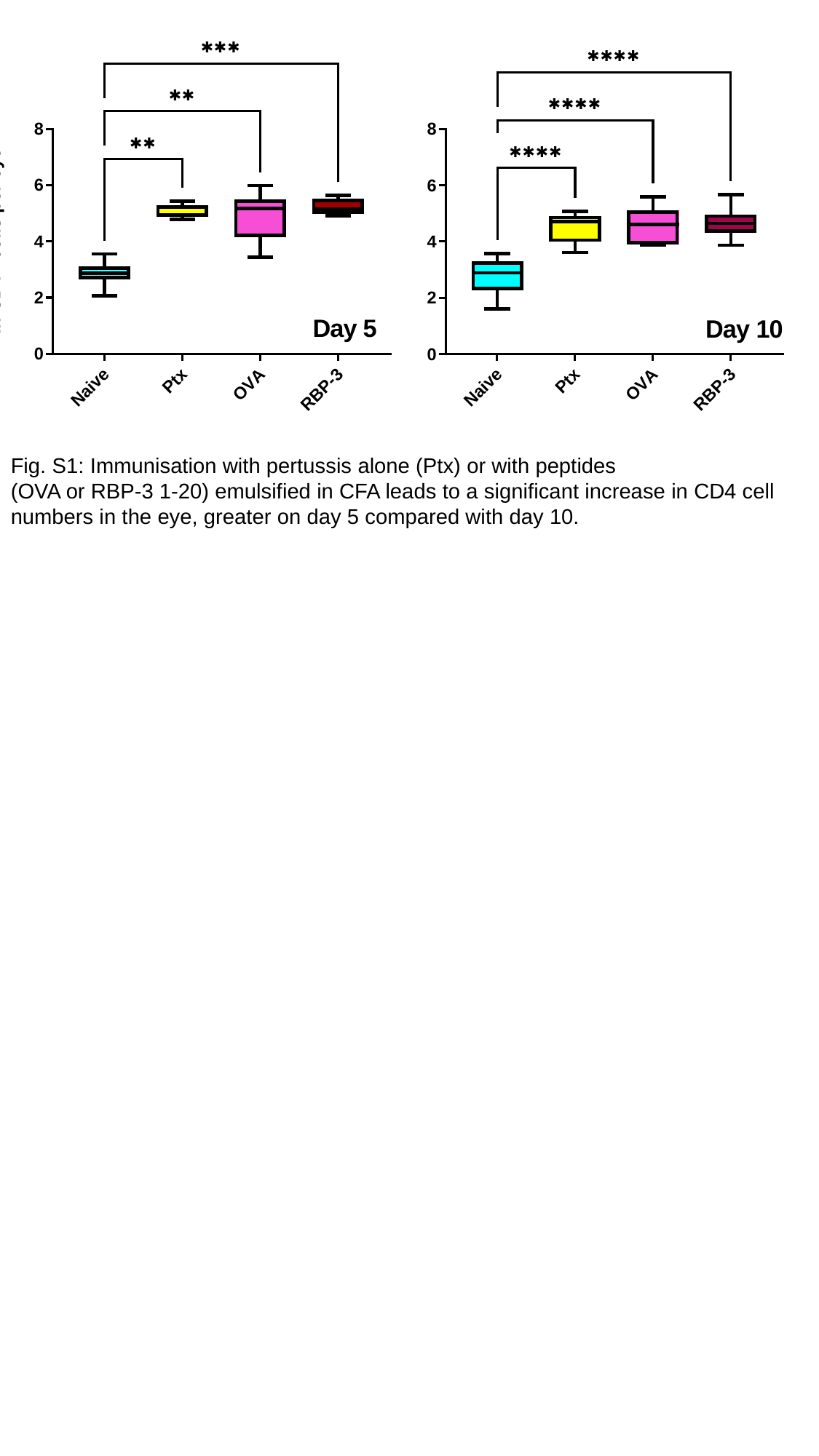

Fig. S1: Immunisation with pertussis alone (Ptx) or with peptides
(OVA or RBP-3 1-20) emulsified in CFA leads to a significant increase in CD4 cell numbers in the eye, greater on day 5 compared with day 10.

### Slide 3
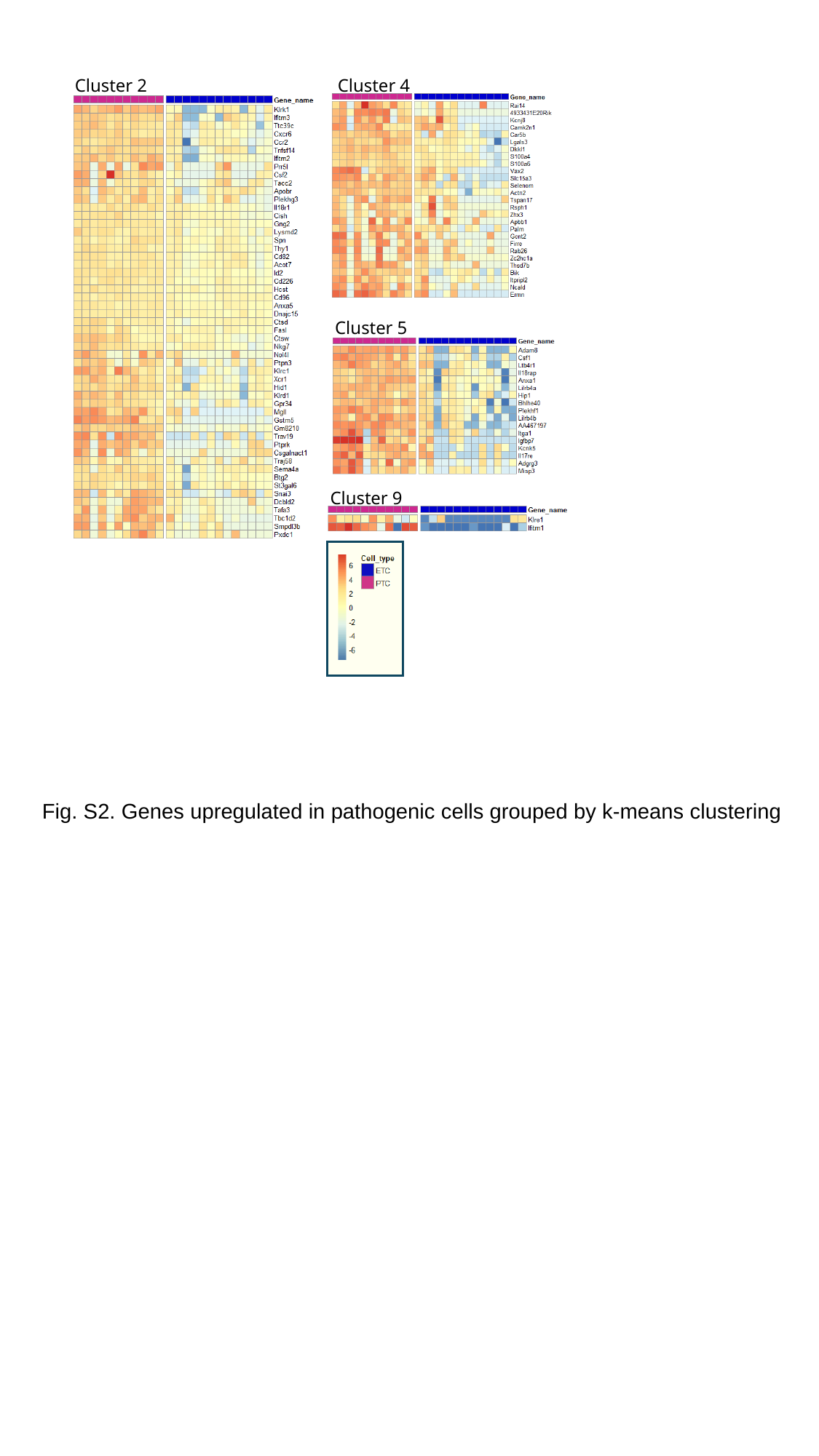

Cluster 2
Cluster 4
Cluster 5
Cluster 9
Fig. S2. Genes upregulated in pathogenic cells grouped by k-means clustering

### Slide 4
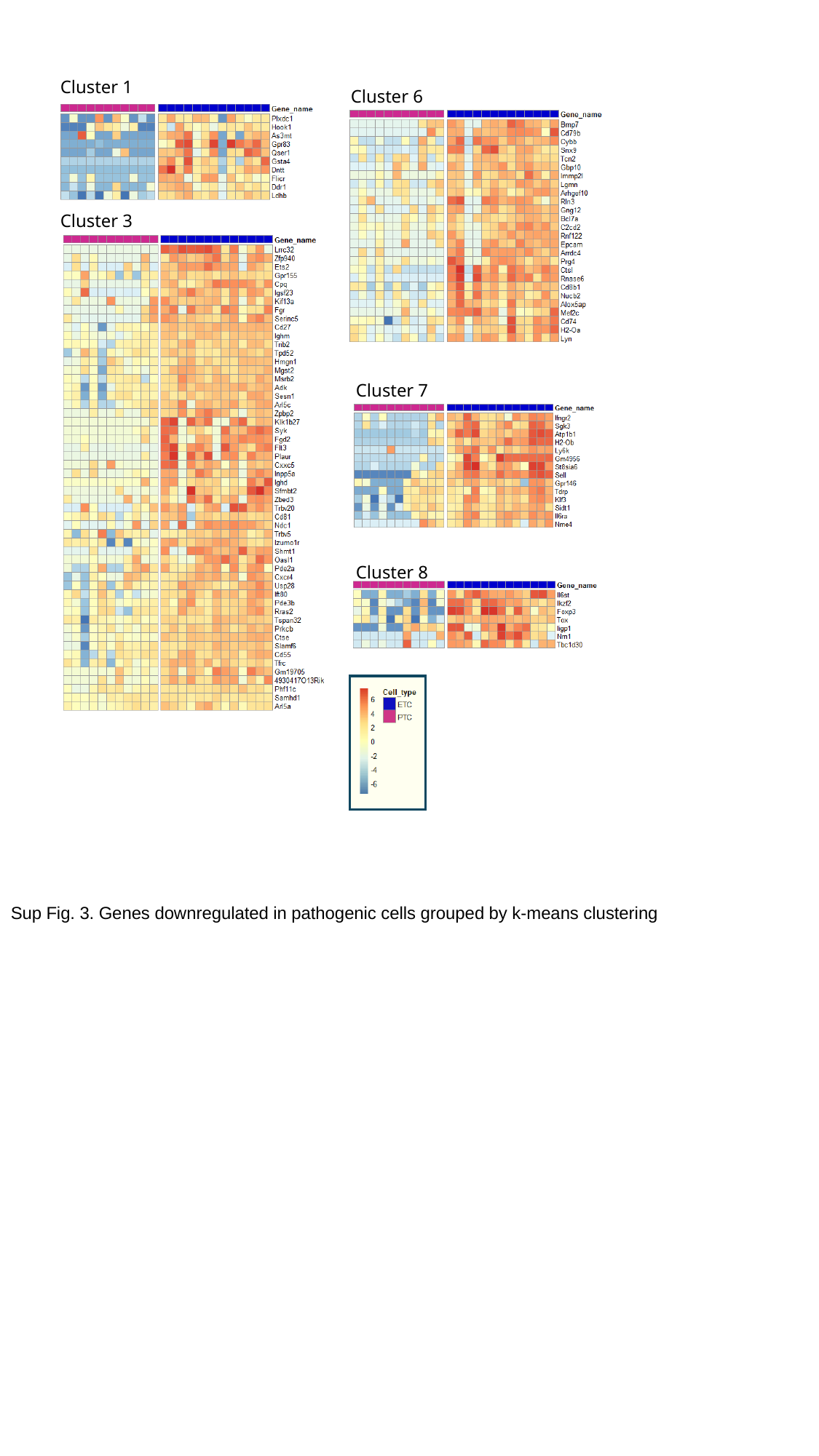

Cluster 1
Cluster 6
Cluster 3
Cluster 7
Cluster 8
Sup Fig. 3. Genes downregulated in pathogenic cells grouped by k-means clustering

### Slide 5
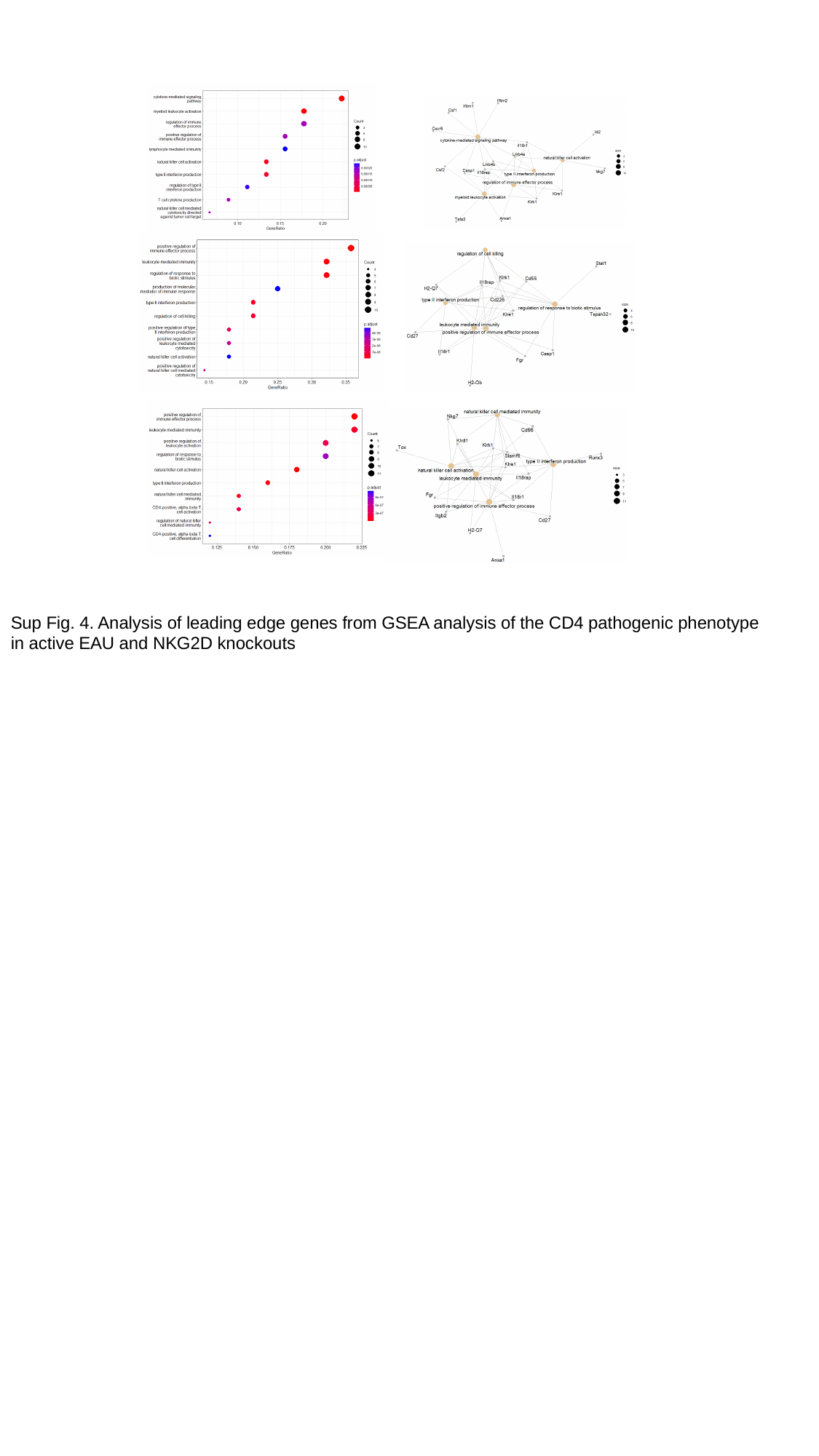

Sup Fig. 4. Analysis of leading edge genes from GSEA analysis of the CD4 pathogenic phenotype in active EAU and NKG2D knockouts
